# The *Trypanosoma brucei* RNA-binding protein DRBD18 ensures correct mRNA *trans* splicing and polyadenylation patterns

**DOI:** 10.1101/2022.03.05.483099

**Authors:** Tania Bishola Tshitenge, Christine Clayton

## Abstract

The parasite *Trypanosoma brucei* grows as bloodstream forms in mammals, and as procyclic forms in tsetse flies. Transcription is polycistronic, all mRNAs are *trans* spliced, and polyadenylation sites are defined by downstream splicing signals. Expression regulation therefore depends heavily on post-transcriptional mechanisms. The RNA-binding protein DRBD18 was previously implicated in the export of some mRNAs from the nucleus in procyclic forms. It copurifies the outer ring of the nuclear pore, mRNA export factors and exon-junction-complex proteins. We show that for >200 mRNAs, DRBD18 depletion caused preferential accumulation of versions with shortened 3’-untranslated regions, arising from use of polyadenylation sites that were either undetectable or rarely seen in non-depleted cells. The shortened mRNAs were often, but not always, more abundant in depleted cells than the corresponding longer versions in normal cells. Their appearance was linked to the appearance of *trans* spliced, polyadenylated RNAs containing only downstream 3’-untranslated-region-derived sequences. Experiments with one mRNA suggested that nuclear retention alone, through depletion of MEX67, did not affect mRNA length, suggesting a specific effect of DRBD18 on processing. Since DRBD18-bound mRNAs were enriched in polypyrimidine tract motifs, and it is found in both the nucleus and the cytoplasm, we suggest that DRBD18 acts in the nucleus by binding to polypyrimidine tracts in 3’-UTRs. DRBD18 binding might both prevent polypyrimidine tract recognition by splicing factors, and promote export of the bound RNAs to the cytosol.

## Introduction

Kinetoplastids are unicellular flagellated parasites that infect mammals and plants. The African trypanosome *Trypanosoma brucei* is a kinetoplastid that causes sleeping sickness in humans in Africa and infects livestock throughout the tropics, with a substantial economic impact (Shaw et al. 2014). *T. brucei* are transmitted by Tsetse flies, or during passive blood transfer by biting flies. In mammalian blood and tissue fluids, the parasites multiply extracellularly as long slender bloodstream forms, escaping the host immune response through antigenic variation of the Variant Surface Glycoproteins (VSGs) (Gray 1965).

In Kinetoplastids, nearly all protein-coding genes are arranged in polycistronic transcription units. Mature mRNAs are generated from the primary transcript by 5’-*trans*-splicing of a 39nt capped leader sequence, and 3’-polyadenylation (Clayton and Michaeli 2011; Michaeli 2011). The parasite regulates mRNAs mainly by post-transcriptional mechanisms, supplemented, in the case of some constitutively abundant mRNAs, by the presence of multiple gene copies. Regulation of mRNA processing, degradation, and translation are therefore central to parasite homeostasis, and for changes in gene expression during differentiation (Fadda et al. 2014; Jensen et al. 2014; Vasquez et al. 2014; Antwi et al. 2016). The sequences required for regulation of mRNA stability and translation often lie in the 3’-untranslated regions (3’-UTRs) of the mRNAs, and most regulation so far has been found to depend on RNA-binding proteins (Clayton 2019). For example, RBP10 (Tb927.8.2780), which is exclusively expressed in mammal-infective forms (Wurst et al. 2012; Savage et al. 2016; Christiano et al. 2017; Shi et al. 2018; Vigneron et al. 2020) specifically associates with procyclic-specific mRNAs, targeting them for destruction (Mugo and Clayton 2017). The *RBP10* 3’-UTR is 7.3kb long, and contains at least six different regions that are independently capable of causing bloodstream-form-specific expression of a reporter (Bishola Tshitenge et al. 2021).

In comparison with mRNA decay and translation, our knowledge of regulatory events during mRNA processing, and especially mRNA export, is rather limited. The mechanism of mRNA *trans*-splicing is basically similar to that for *cis* and *trans* splicing in other eukaryotes (Michaeli 2011; Preusser et al. 2012), as is the polyadenylation machinery (Koch et al. 2016). Unusually, however, the two processes are inextricably linked: the polyadenylation site of each mRNA is determined solely by the location of the next downstream splice site (Huang and van der Ploeg 1991; Ullu et al. 1993; Matthews et al. 1994; Vassella et al. 1994); and chemical or RNAi-mediated inhibition of either splicing or polyadenylation stops both processes (Ullu and Tschudi 1991; McNally and Agabian 1992; Hendriks et al. 2003; Begolo et al. 2018; Wall et al. 2018). Like other eukaryotes, trypanosomes have an exon junction complex, which is presumably deposited at the end of the spliced leader after or during processing: it includes Y14 (Tb927.7.1170), Magoh (Tb927.6.4950) and an NTF2-like protein (Tb927.10.2240) (Bercovich et al. 2009) and perhaps an eIF4AIII homologue, Tb927.11.8770 (Dhalia et al. 2006; Inoue et al. 2014). Bioinformatic analysis and limited reporter experiments indicate that - as in other eukaryotes - a polypyrimidine tract (PPT) serves as the main *trans*-splicing signal (Matthews et al. 1994; Schürch et al. 1994; Vassella et al. 1994; Nilsson et al. 2010; Siegel et al. 2010), but in contrast with at least some eukaryotes (Patzelt et al. 1989), there is no consensus branch-point sequence. The results of reporter experiments indicate that the nature of the sequence downstream of the splice site, as well as the length and composition of the PPT, influence splice site choice (Hartmann et al. 1998; Lopez-Estrano et al. 1998; Siegel et al. 2005). There are also indications that some mRNAs are more efficiently spliced than others (Kapotas and Bellofatto 1993; Fadda et al. 2014; Antwi et al. 2016); and when processing is inhibited, the precursors are substrates for the exosome (Kramer et al. 2016). Several RNA-binding proteins have been shown to influence splicing (Gupta et al. 2013a; Gupta et al. 2013b; Gupta et al. 2014) but their mechanisms of action are not clear; often, they have double functions, also binding to the 3’-UTRs of mature mRNAs and affecting mRNA stability.

The mechanism of mRNA export from the nucleus in Kinetoplastids has received relatively little attention (reviewed in (Kramer 2021)). In Opisthokonts, mRNA export requires a complex that is conserved across eukaryotes: it includes Mex67 (NXF1 or TAP in mammals) and the helicase Mtr2 (p15 or NXT1 in mammals) as well as various other less conserved proteins (reviewed in (Ashkenazy-Titelman et al. 2020)). Mex67 interacts with both the mRNA and the nuclear pore. Gle2 (RAE1), a nuclear pore component, is also implicated in export. Recognition of mature mRNAs by export factors involves various proteins that can bind to the RNA, including cap-binding proteins, splicing factors and the exon junction complex (reviewed in (Ashkenazy-Titelman et al. 2020)). Trypanosomes use a complex that contains the major export factor MEX67 (Tb927.11.2370) (Schwede et al. 2009; Dostalova et al. 2013), the helicase MTR2 (Tb927.7.5760), and a transportin-like protein, IMP1 (Tb927.9.13520) (Dostalova et al. 2013); trypanosome GLE2 (Tb927.9.3760) is also implicated. The inner ring of the trypanosome nuclear pore is very similar to those of Opisthokonts, but outer components are more diverged and there are no cytoplasmic filaments (Obado et al. 2016). In trypanosomes export of mRNAs can be initiated before they have been completely transcribed (Goos et al. 2018), so recognition must depend on the 5’-end: we do not know whether the export machinery recognises the cap, cap-binding proteins, or the exon junction complex. The TREX and THO complexes, which are implicated in the integration of transcription, splicing and export in Opisthokonts (Ashkenazy-Titelman et al. 2020), are absent in trypanosomes (Kramer 2021).

The double RNA-binding protein 18 (DRBD18) was initially identified as a substrate of arginine methylation (Lott et al. 2015). It is expressed in both bloodstream and procyclic forms. Various lines of evidence indicate that DRBD18 is involved in the export of a subset of mRNAs from the nucleus. DRBD18 interacts directly with MTR2, and co-immunoprecipitates MEX67 (Mishra et al. 2021). Moreover, in procyclic forms, depletion of DRBD18 by RNAi caused partial retention of MTR2 and MEX67, and some poly(A)+ mRNAs, in the nucleus (Mishra et al. 2021). Levels of various mRNAs were affected, but the effects were different depending on whether cytoplasmic or whole cell mRNAs were analysed (Mishra et al. 2021).

One of the mRNAs that increased significantly in abundance after DRBD18 depletion in procyclic forms was that encoding RBP10 (Lott et al. 2015; Mishra et al. 2021). During studies of RBP10 regulation, we unexpectedly discovered that DRBD18 depletion led to accumulation of truncated versions of the *RBP10* mRNA. Analysis of transcriptomes from both bloodstream and procyclic forms revealed that DRBD18 depletion affects processing of over 200 mRNAs. These results suggest that DRBD18 may affect mRNA processing as well as mRNA export.

## Results

### DRBD18 depletion affects *RBP10* mRNA processing

We initially studied DRBD18 because its depletion in procyclic forms had been shown to cause a modest increase in the abundance of *RBP10* mRNA (Lott et al. 2015; Mishra et al. 2021). We found that depletion of DRBD18 in bloodstream forms caused growth inhibition (Figure 1A) and a modest (3-fold) decrease in RBP10 protein (Figure 1B). Remarkably, however, when we looked at *RBP10* mRNA by Northern blotting, we discovered that DRBD18 depletion caused accumulation of progressively smaller *RBP10* mRNAs (Figure 1C). Since the blot was hybridised to the open reading frame (coding sequence, CDS) probe, the mRNAs must have different 3’-UTRs. Eukaryotic mRNAs that lack poly(A) tails are generally extremely unstable, so we hypothesised that the various *RBP10* mRNAs were products of alternative polyadenylation. To find out whether the effect was specific for *RBP10*, we examined mRNA from the tandemly repeated tubulin genes, because processing inhibition causes accumulation of tubulin (*TUB*) RNA dimers and multimers. Processing of *TUB* mRNA was not affected, showing that the effect on *RBP10* mRNA was not due to a general RNA processing defect (Figure 1C). A similar effect on *RBP10* mRNA was seen after DRBD18 depletion in a different strain, EATRO1125 (Supplementary Figure S1A, B). Analysis of 3’-ends of the different *RBP10* mRNAs by RT-PCR (3’-RACE) using oligo (dT)_18_ was not successful, perhaps because the sequence is generally of low complexity and the forward primers chosen were similar to sequences in other mRNAs.

**Figure 1:**
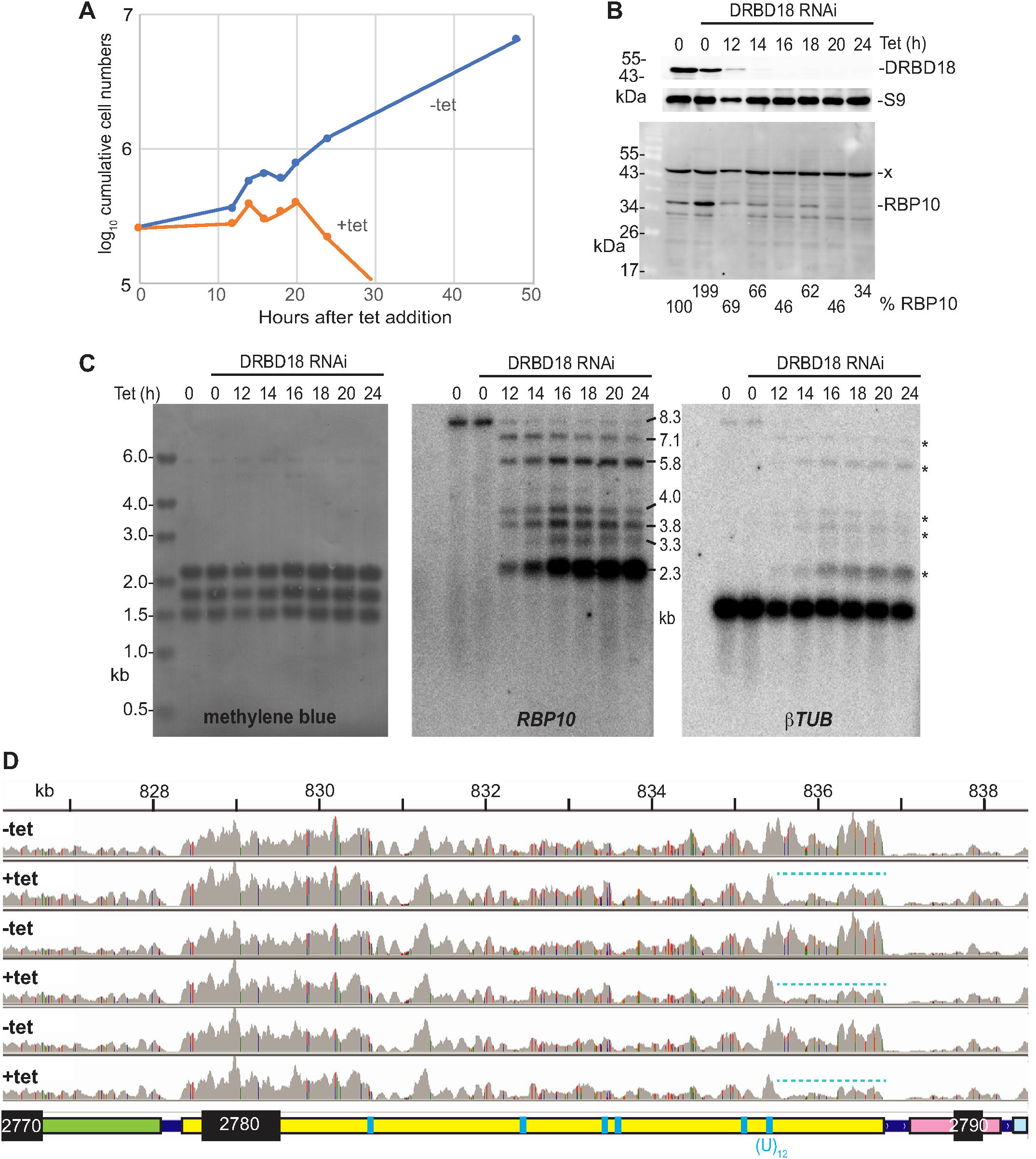
Effect of *DRBD18* RNAi on *RBP10* mRNA and expression levels. **A.** Example of cumulative growth curves of bloodstream forms with and without induction of *DRBD18* RNAi. **B.** DRBD18 and RBP10 protein levels after RNAi induction, corresponding to (A). Ribosomal protein S9 is the loading control. Anti-RBP10 antibodies (Wurst et al. 2012) were used for detection of RBP10 and a non-specific band (x). **C.** RNA from the samples used in (A) was used to examine effects on *RBP10* mRNA. The methylene blue-stained membrane is shown on the left, with *RBP10* hybridisation in the centre. The membrane was then stripped and hybridised with a beta-tubulin probe as a further control; *\*RBP10* signal that remained after stripping. **D.** Visualisation of RNA-Seq results after 12h tetracycline-mediated induction of *DRBD18* RNAi, using the Integrated Genomics Viewer (Robinson et al. 2011; Thorvaldsdóttir et al. 2013). Three replicates are shown. Note that these results are shown on a linear scale whereas those in Figure 1A are on a log scale. Proposed full length mRNAs are shown below the tracks; positions of (U)_12_ or (U)_6_C(U)_6_ sequences are indicated in cyan but there are numerous other polypyrimidine tracts present. The cyan dotted line highlights loss of reads at the end of the *RBP10* 3’-UTR after 12h *DRBD18* RNAi. The read densities over the middle portion of the 3’-UTR are lower than the rest, probably because some of the reads will be aligned to the additional sequence copy in the TREU927 reference genome. Coloured lines indicate read mismatches with the TREU927 reference genome.

To examine this in more detail and to look for effects on other mRNAs, we examined the transcriptomes of bloodstream forms 12 hours after DRBD18 RNAi induction. This time was chosen as the first point at which DRBD18 was completely depleted without much effect on growth (Figure 1A,B). We sequenced total rRNA-depleted RNA. The numbers of reads from the *RBP10* coding region were not much affected after 12h (Supplementary Table S1), as expected from the Northern blot results. Visualisation in the Integrated Genome Viewer (Robinson et al. 2011; Thorvaldsdóttir et al. 2013) also showed that with or without tetracycline induction of RNAi, the read density over the *RBP10* coding region was 2-3 times higher than the density over the upstream gene, Tb927.8.2770 (Figure 1D). However, over the last 2kb of the 3’-UTR, there were far fewer reads after DRBD18 depletion (cyan dotted line), indicating loss of the longest *RBP10* mRNAs and consistent with the Northern blot result.

The results so far suggested that multiple alternative polyadenylation sites were being used after *DRBD18* depletion. The *RBP10* 3’-UTR contains several PPTs that might serve as splicing signals (Figure 1D).

### Depletion of MEX67 does not cause accumulation of shortened *RBP10* mRNAs

A previous paper implicated DRBD18 in export of mRNAs from the nucleus. We had previously shown that induction of RNAi targeting *MEX67* caused accumulation of poly(A)+ RNA in the nucleus (Schwede et al. 2009). After 24h RNAi induction in this cell line, growth of the parasites was clearly inhibited (Figure 2A) and *MEX67* mRNA was reduced (Figure 2B), but the migration of *RBP10* mRNA was unaffected (Figure 2B). This suggests that altered processing of *RBP10* mRNA after DRBD18 depletion was not caused by retention in the nucleus alone.

**Figure 2:**
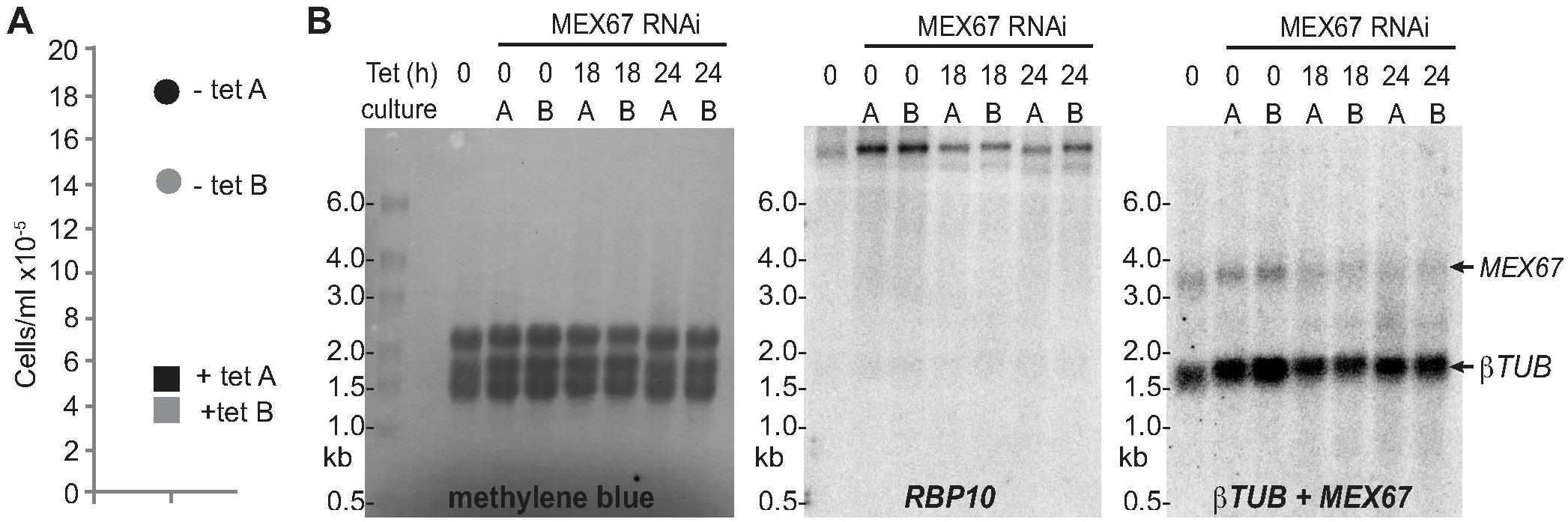
Effect of *MEX67* RNAi *on RBP10* mRNA and expression levels. **A.** Cell densities in two cultures 24h after addition of tetracycline to induce *MEX67* RNAi. **B.** Northern blots using RNA from the two cultures, with detection of rRNA (methylene blue) and of *RBP10* and beta-tubulin mRNAs.

### Composition and associations of proteins of DRBD18 mRNPs

There are two lines of evidence that DRBD18 is involved in mRNA export. The strongest argument is that its depletion in procyclic forms causes retention of mRNAs in the nucleus (Mishra et al. 2021). The second type of evidence is from protein-protein interactions. An RNA-independent interaction with MTR2 and MEX67 was confirmed by co-immunoprecipitation in the presence of RNase (Mishra et al. 2021) and a two-hybrid interaction with MTR2 was also demonstrated. In addition, non-quantitative mass spectrometry results with DRBD18 preparations from procyclic forms suggested co-purification not only with MEX67 and MTR2, but also some nuclear pore components and various RNA-binding proteins (Lott et al. 2015). Since label-free quantitative mass spectrometry is now readily available, we used it to re-examine DRBD18 protein associations in more detail. As some direct interactions of DRBD18 had already been demonstrated, we decided to focus on the components of DRBD18-containing “messenger ribonucleoprotein particles” (mRNPs) and their interactions, including other proteins that might be bound via RNA. DRBD18 with a C-terminal TAP (tandem affinity purification) tag was expressed from one allele in bloodstream forms. We do not know whether this protein was fully functional, since attempts to delete the other (untagged) copy failed. It was not possible to obtain cells expressing DRBD18 with N-terminal tags. We used just a single step of purification in order to avoid loss of material and interaction partners. DRBD18-TAP was pulled-down using IgG beads, then released by Tobacco Etch Virus (TEV) protease cleavage at a site within the tag. The resulting mixture was analysed by quantitative mass spectrometry (Supplementary Table S2). GFP-TAP and TAP-ZC3H28 (Bishola Tshitenge and Clayton 2021) served as controls. ZC3H28 is an RNA-binding protein that is predominantly in the cytoplasm (Bishola Tshitenge and Clayton 2021).

Figure 3 shows analysis of the mass spectrometry data using the Perseus algorithm (Tyanova et al. 2016). The major advantage of Perseus is that it yields probability values that are adjusted for multiple testing. A disadvantage is that it handles absent values by substituting simulated intensities, which can lead to false negative results if a protein that interacts with the investigated protein is not detected in the negative control. In the following discussion we consider two overlapping sets of proteins as being significantly enriched: those that had adjusted P-values of less than 0.01, with greater than 4-fold enrichment as calculated in Perseus, and those that were detected with at least one peptide in all of the DRBD18 preparations but in none of the control preparations (Supplementary Table S2). The comparisons showed that DRBD18-TAP specifically copurified several sets of proteins. We found MTR2 as expected, but also other proteins linked to mRNA export from the nucleus: a transportin-2-like protein, an importin-beta subunit, a Ran binding protein, the putative nuclear RNA helicase EIF4AIII and a possible Ran-GAP. EIF4AIII was however also in two of the three ZC3H28 preparations. MEX67 was found in two of the three DRBD18 preparations and none of the others, but GLE2 was not detected. The exon junction complex components Magoh and NTF2-domain-protein (Bercovich et al. 2009) were found with both DRBD18 and ZC3H28, but not the GFP control, but Magoh was more reproducibly associated with DRBD18. Most dramatically, all components of the outer ring of the nuclear pore (Degrasse et al. 2009; Obado et al. 2016; Obado et al. 2017) purified highly specifically with DRBD18. All of these results are consistent with a role of DRBD18 in mRNA export.

**Figure 3:**
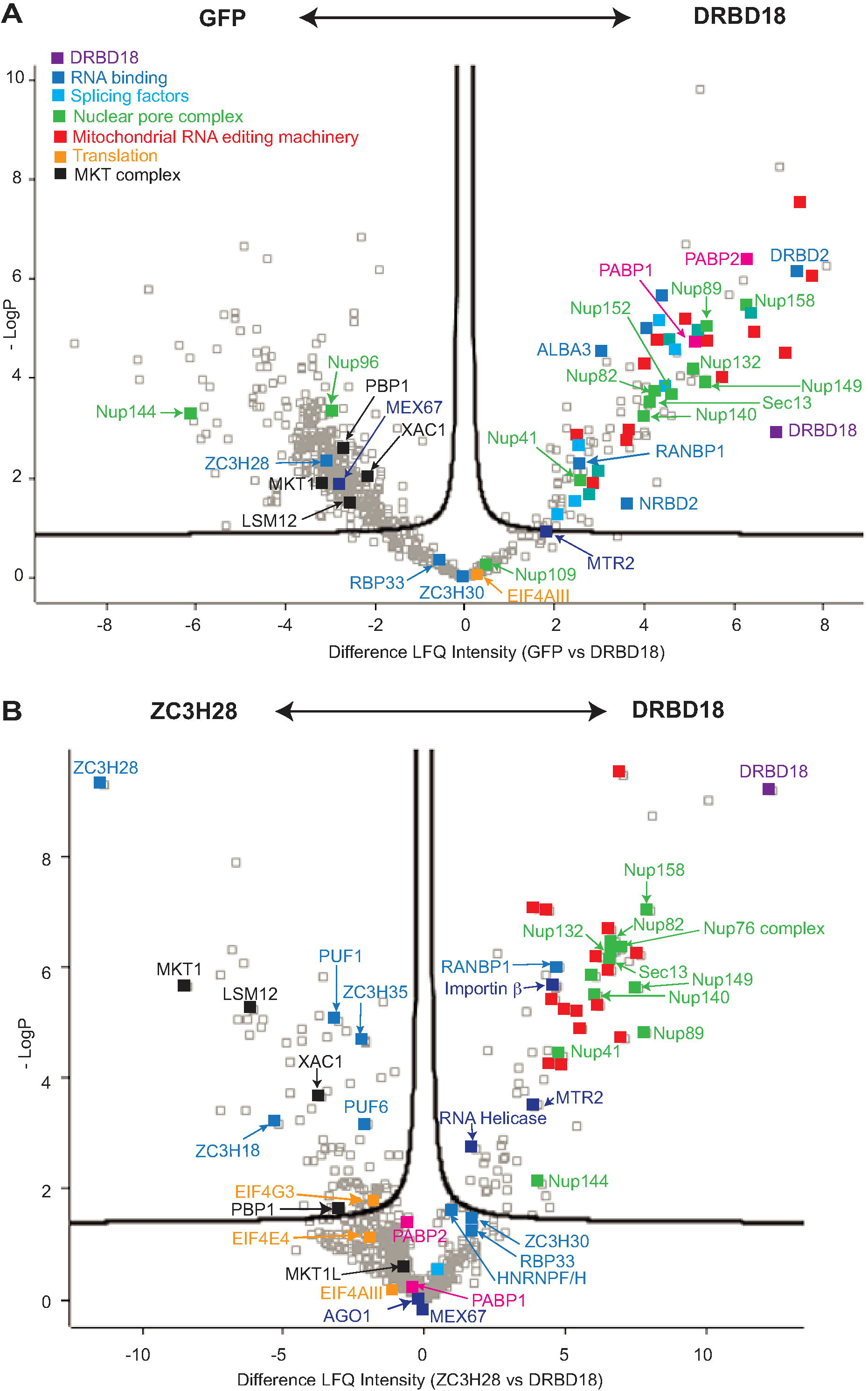
Proteins associated with DRBD18. **A.** Volcano plot showing proteins that were significantly enriched with DRBD18-TAP relative to TAP-GFP. Proteins of interest are labelled. The x-axis shows log_2_ enrichment, while the y-axis is -log_10_ of the P-value generated by the Perseus algorithm (Tyanova et al. 2016). Proteins outside the curved line are significantly enriched. **B.** Volcano plot showing proteins that were significantly enriched with DRBD18-TAP relative to TAP-ZC3H28. The data for ZC3H28 and GFP were described previously (Bishola Tshitenge and Clayton 2021).

The mass spectrometry results also gave some indications of other proteins that are enriched in DRBD18-containing mRNPs. The RNA-binding domain proteins ZFP2, RBP33 and ZC3H30 were specifically associated with DRBD18, and not ZC3H28 mRNPs. ZFP2 is required for differentiation of bloodstream to procyclic forms (Hendriks et al. 2001). The function of ZC3H30 is unknown (Chakraborty and Clayton 2018). RBP33 is an essential nuclear protein, so its association with DRBD18 but not ZC3H28 corresponds to the localisations of the two proteins. RBP33 associates with non-coding RNAs, especially from antisense transcription (Fernandez-Moya et al. 2014), but its depletion led to loss of mRNA and spliced leader RNA (Cirovic et al. 2017), suggesting a link to splicing. Both DRBD18 and ZC3H28 enriched various ribosomal proteins and RNA-binding proteins, including both poly(A) binding proteins (PABP1 and PABP2), ALBA2, ALBA3, DRBD2, DRBD3, ZC3H9, ZC3H34, ZC3H39, ZC3H40, UPF1, HNRNPH/F and TRRM1. There were also a few splicing factors, the RNA interference Argonaute protein AGO1, and the translation initiation complex EIF4E3/4G4 (Supplementary Table S2). Many of these proteins are likely to associate via RNA. Unlike ZC3H28, DRBD18 was not associated with MKT1, LSM12, XAC1, and PBP1, which are components of a complex that promotes translation and mRNA stability in the cytoplasm (Singh et al. 2014; Nascimento et al. 2020). Association of PUF1, PUF6, ZC3H35, and ZC3H18 was also specific to ZC3H28. Intriguingly, DRBD18 specifically copurified multiple components of the mitochondrial RNA editing machinery; this might warrant future investigation.

We next checked the location of DRBD18, since existing results were contradictory: Lott *et al*. (Lott et al. 2015) reported that it was in the cytosol whereas C-terminally GFP-tagged DRBD18 was in the nucleus (Dean et al. 2016). After cell fractionation, the control cytoplasmic protein RBP10 and the nuclear exoribonuclease XRND (Li et al. 2006) showed the expected distributions. In contrast, we found that DRBD18 was associated with both compartments (Supplementary Figure S1C, D), which is consistent with a role in export of mRNAs from the nucleus.

### Long *RBP10* transcripts are mainly in the nucleus after DRBD18 depletion, while shorter ones are exported

Our mass spectrometry results (Figure 3) supported previous observations implicating DRBD18 in mRNA export. Inhibition of export alone was however not sufficient to cause accumulation of shorter *RBP10* mRNA variants (Figure 2), and the previous publication had already indicated that DRBD18 depletion affects the distributions of only some mRNAs (Mishra et al. 2021). We hypothesised, therefore, that DRBD18 binds to possible processing sites in the *RBP10* 3’-UTR, and that this not only promotes efficient export of the long mRNA, but also inhibits the use of the alternative sites. If this were true, DRBD18 RNAi might prevent export of longer *RBP10* mRNAs, while simultaneously allowing them to be processed to shorter versions. To test this, we examined the locations of the different *RBP10* mRNAs by single-molecule fluorescent *in situ* hybridization. To distinguish the different mRNAs, we used four probes (Figure 4A). Details of the results are in Supplementary Table S3. The shortest species detected had signals from the coding region only (pink probe 1) and were mostly nuclear: these could represent precursors that were still undergoing transcription. The next shortest mRNAs detected hybridised only to probes 1 (coding region) and 2. These must be 2-7 kb long (Figure 4A, B). The mRNAs that hybridise to probes 1 and 3 are at least of intermediate (or medium) length (5-7 kb long); and those that hybridise to probes 1,2 and 4 or 1,3 and 4 are more than 7.5 kb long (Figure 4A, B). On average, 5% of the mRNAs hybridised only to probes 1 and 4: this is physically unlikely and may indicate of the background failure rate for hybridisation of probes 2 or 3 (Supplementary Table S3). The numbers of mRNAs hybridising to probes 2 or 3 only were unexpectedly high (average 17% of signals); although these do not match any other sequences in the Lister427 genome, they do contain many low complexity sequences which might result in cross-hybridisation. RNAs that hybridised to the green probe only, or to the red and green probes, could be artefacts, or represent products either from degradation, or from alternative splicing using 3’-UTR-internal signals (Kramer 2017). There were very few of either and the numbers were not reproducibly affected by *DRBD18* RNAi. If all mRNAs hybridising to the coding region probe were included, wild-type cells contained 3-3.5 *RBP10* mRNAs per cell; this is consistent with previous calculations (Fadda et al. 2014), which suggested 4 *RBP10* mRNAs per cell.

**Figure 4:**
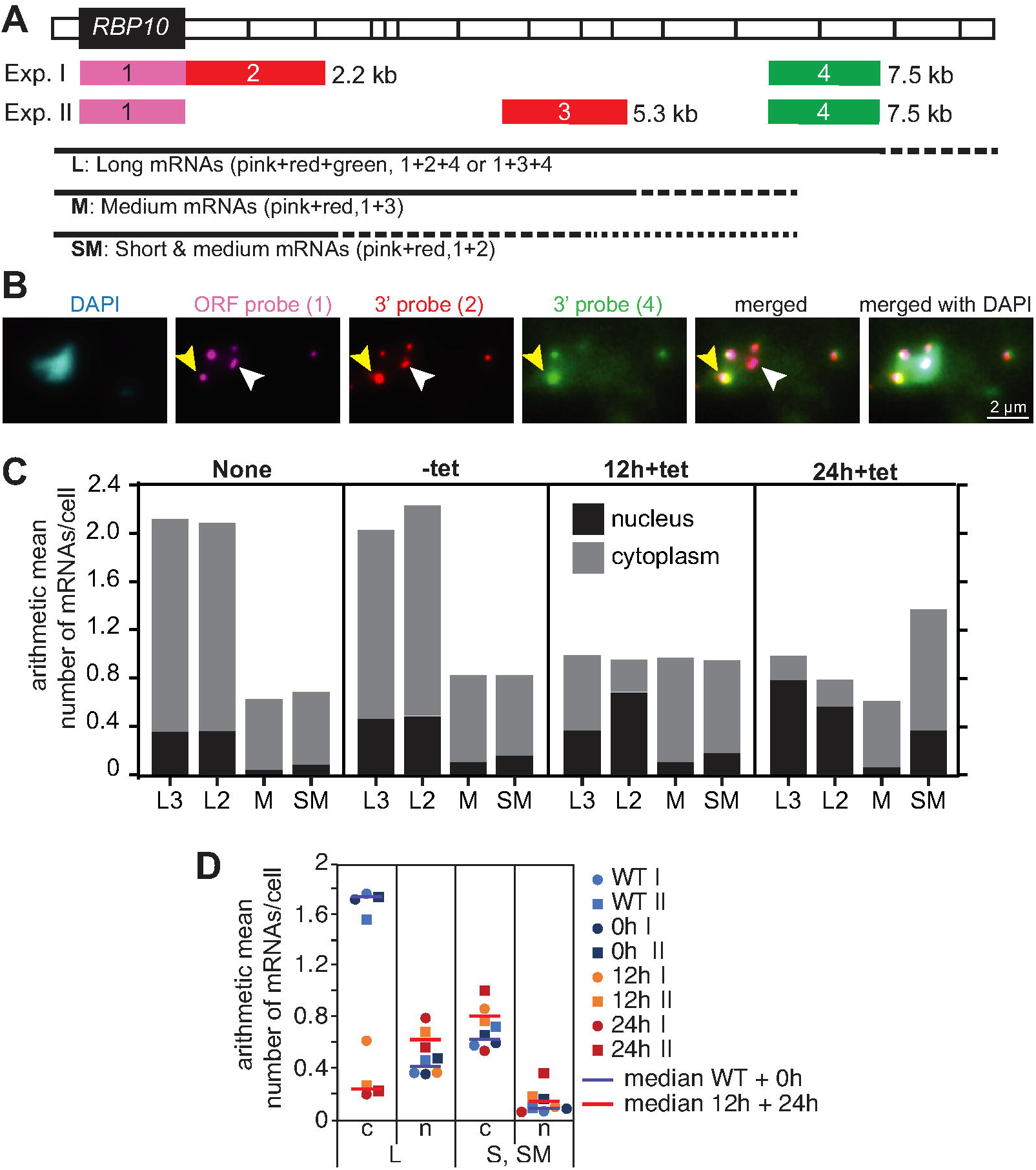
Long *RBP10* mRNAs remain in the nucleus after DRBD18 depletion. Individual *RBP10* mRNAs were detected by single-molecule fluorescent *in situ* hybridization with or without DRBD18 depletion for 12 or 24h. In each case, 50 cells were analysed. **A.** Map showing the positions of the probes on the *RBP10* mRNA, approximately to scale. The 3’-UTR divisions on the map correspond to those in Figure 2. The bars indicate the extent of mRNAs detected by the probe combinations. The dotted lines indicate that RNAs hybridising to the different probe sets could have various lengths. The distances next to the bars indicate the position of the probe end on the mRNA. Exp.: experiment. **B.** Example image of *RBP10* mRNAs in a wild-type cell, detected by single-molecule fluorescent *in situ* hybridization using probes 1,2 and 4 (experiment 1 in panel (A)). DNA was stained with DAPI. The white arrow indicates an mRNA that hybridises only to the coding region (open reading frame, ORF) and 3’ probe 2; the yellow arrow points to one of the four mRNAs that hybridise to probes 1,2 and 4. **C.** Mean numbers of long, medium and short mRNAs per cell under different conditions. The data were from 50 cells. “None”: Cells with no RNAi plasmid; “-tet”: RNAi cells without tetracycline; and cells after 12h or 24h tetracycline treatment. All slides were hybridised with the red, pink and green probes, and results for hybridisation with the two different red probes are shown separately. “L3” indicates long mRNAs that hybridised with probes 1,3 and 4 (experiment II) and “L2” indicates long mRNAs that hybridised with probes 1,2 and 4 (experiment I). The black bars indicate mRNAs in the nucleus and the grey bars, mRNAs in the cytoplasm. Numbers for mRNAs hybridising to only one probe, or red and green only, are in Supplementary Table S3. **D.** Compilation of data from panel C, to allow comparison of the numbers of full-length and shorter mRNAs in the nucleus and cytoplasm. The key to symbols is on the right.

In order to be certain that only specific signals are considered, we decided to consider only the long, medium and short mRNAs (as defined above) which hybridised to at least 2 probes. In wild-type cells, about 70% of *RBP10* mRNAs were longer than 7.5 kb; 17% of the long mRNAs and 9-12% of the short or medium ones were in the nucleus (Figure 4C). Cells containing the *DRBD18* RNAi plasmid, but with no induction, showed similar values to wild-type (Figure 4C,D). After RNAi induction, the total number of fulllength *RBP10* mRNAs was roughly halved (Student t-test p-value 2 x 10^-6^) (Figure 4D) and the number in the cytoplasm had decreased 7-fold (Student t-test p-value <.0002). Thus most of the full-length mRNAs that remained were now in the nucleus (Figure 4C,D). (Although the median number in the nucleus had increased about 1.5-fold, the difference was not statistically significant.) This mirrored the Northern blot results. The amounts of medium-length and shorter *RBP10* mRNAs without RNAi were higher than expected from the Northern blots, while the 1.4-fold increase after RNAi was unexpectedly small. However, the short and medium mRNAs were more than 4-fold more abundant in the cytoplasm than in the nucleus at most time points, indicating that their export was unaffected by DRBD18 depletion. We know that reporter mRNAs with shortened *RBP10* 3’-UTRs can be translated (Bishola Tshitenge et al. 2021), so the reduced amount of RBP10 protein after DRBD18 RNAi (Figure 1B) is probably caused by the decrease in total cytosolic *RBP10* mRNA (Figure 4C,D).

### Effects of DRBD18 depletion on mRNA abundance

DRBD18 depletion affected the length of *RBP10* mRNA, but not tubulin mRNAs. To find out whether *DRBD18* RNAi affects processing of additional mRNAs, we first examined the transcriptomes of *DRBD18*-depleted bloodstream forms (Supplementary Table S1) manually using the Integrative Genomics Viewer (Robinson et al. 2011), focussing on mRNAs with long 3’-UTRs. We found several that appeared to have altered processing. Northern blotting confirmed that one example, *DRBD12* mRNA - which is usually about 8.4 kb long - accumulated as a 2.6 kb mRNA from 12h *DRBD18* RNAi onwards, with an intermediate band of about 4kb (Supplementary Figure S2). We therefore undertook more detailed analysis of all available datasets. In addition to our bloodstream-form transcriptomes (12h RNAi, whole-cell total RNA, processed with oligonucleotides and RNase H to deplete rRNA) (Supplementary Table S1), we examined the transcriptome data from procyclic forms after 19h RNAi (Mishra et al. 2021). The procyclic-form RNA was poly(A)+ mRNA purified, so we could be certain that it was processed; moreover, results are available both from whole cells and from the cytoplasmic fraction (Supplementary Table S4). Transcripts that are more abundant in the whole-cell mRNA than in the cytoplasmic mRNA are either poorly exported, associated with the nuclear membrane, or unstable in the cytoplasm.

We first looked at effects on mRNA abundance using the reads from coding regions, using DeSeq2 (Love et al. 2014; Leiss and Clayton 2016). There was only weak correlation between the effects in bloodstream and procyclic forms (Pearson correlation coefficient 0.32) (Supplementary Figure S3A). However, considering only the significantly affected genes (adjusted P-value <0.05, change of at least 1.5-fold) there was considerable overlap. Using a set of “unique” genes that excludes paralogues (Supplementary Table S5, sheet 4), and also excluding *DRBD18* itself, 397 mRNAs were decreased in bloodstream forms, and 81 in procyclic forms; of these, 61 were decreased in both (Supplementary Table S5 sheet 4) (Fisher test P=4 x 10^-60^). There were no enriched funtional categories among the mRNAs that were decreased in both forms.186 mRNAs were 1.5-fold significantly (Padj <0.05) increased in bloodstream forms and 304 in procyclic forms; of these, 72 were increased in both (Supplementary Table S5 sheet 4) (Fisher test P=4×10^-51^).

In bloodstream forms, a minority of the RNA abundance changes after DRBD18 RNAi may have been due to the onset of growth arrest, or to stress. Examples include decreases in mRNAs encoding some cytoskeletal proteins, enzymes of glucose or glycerol metabolism, and increases in mRNAs encoding membrane- and mitochondrial proteins; such mRNAs also showed equivalent changes in the growth-arrested stumpy form (Naguleswaran et al. 2018; Silvester et al. 2018; Quintana et al. 2021). For the procyclic forms there was no significant overlap between the effects of DRBD18 depletion and the transcriptomes of growth-arrested metacyclic forms (Christiano et al. 2017). There was also no correlation between effects of DRBD18 depletion and differences between wild-type bloodstream and procyclic forms (not shown).

Among the 72 mRNAs that were increased in both stages after RNAi, 9 encoded RNA-binding proteins. This represented significant enrichment (adjusted Fisher P-value = 3×10^-6^). However, these mRNAs have long 3’-UTRs. For seven of the nine, there was evidence for truncation of the 3’-UTR after DRBD18 RNAi (Supplementary Table S7, sheet 2). In other cases, particularly for the mRNAs affected only in procyclic forms, the read coverage revealed no evidence for changes in processing, suggesting that DRBD18 loss had affected mRNA stability.

### DRBD18 is associated with mRNAs that have long 3’-UTRs containing polypyrimidine tracts

To find out whether mRNAs that were affected by *DRBD18* RNAi were also bound by DRBD18 at steady state, we purified DRBD18-TAP, released the protein with TEV protease, and sequenced the both the eluted RNAs and those that did not bind to the beads (Supplementary Table S7). We found 226 mRNAs that were at least 2-fold enriched (normalised elute reads divided by corresponding unbound reads) in all three replicates, and defined these as “bound” RNAs (Supplementary Table S7, sheet 1). There was no overall correlation between binding of mature transcript by DRBD18 and effects on transcript abundance (Supplementary Figure S3B), and no correlation with retention in the nucleus after DRBD18 depletion (Mishra et al. 2021) (Supplementary Table S4, sheet 5 and Supplementary Table S7, sheet 1). Of the 36 bound mRNAs that were at least 1.5-fold changed by DRBD18 depletion, 25 increased (Fisher p-value 4 x 10^-10^) whereas 10 decreased (Fisher p-value 0.9). At least three of the bound and increased mRNAs clearly had truncated 3’-UTRs (see below). *RBP10* mRNA was more than 2-fold enriched in two of the three replicates.

Binding of DRBD18 did not correlate overall with either mRNA (Supplementary Figure S3C) or 3’-UTR length (Figure 5A). However, the 3’-UTRs of the “bound” mRNAs were significantly longer than those of mRNAs that were, on average, less than one-fold enriched (“unbound”) (Figure 5A, B). Lott et al (Lott et al. 2015) found a poly(A)-rich motif in mRNAs that were decreased after DRBD18 RNAi, and CACCCAC motif in mRNAs that increased after DRBD18 RNAi. The functions of those motifs were however not tested for roles in either the RNAi response or DRBD18 binding. When the 3’-UTRs of our “bound” mRNAs were compared with length-matched controls (Supplementary Table S6 sheet 3) using MEME (Bailey et al. 2015), in the discriminative mode and looking for 6-20nt motifs, three sequences were found (Figure 5C). Of these, the one that was present in the most mRNAs (157/163) was a PPT. Only two motifs were identified by MEME in the differential enrichment mode, with lower significance, but they were again PPTs (Figure 5C).

**Figure 5:**
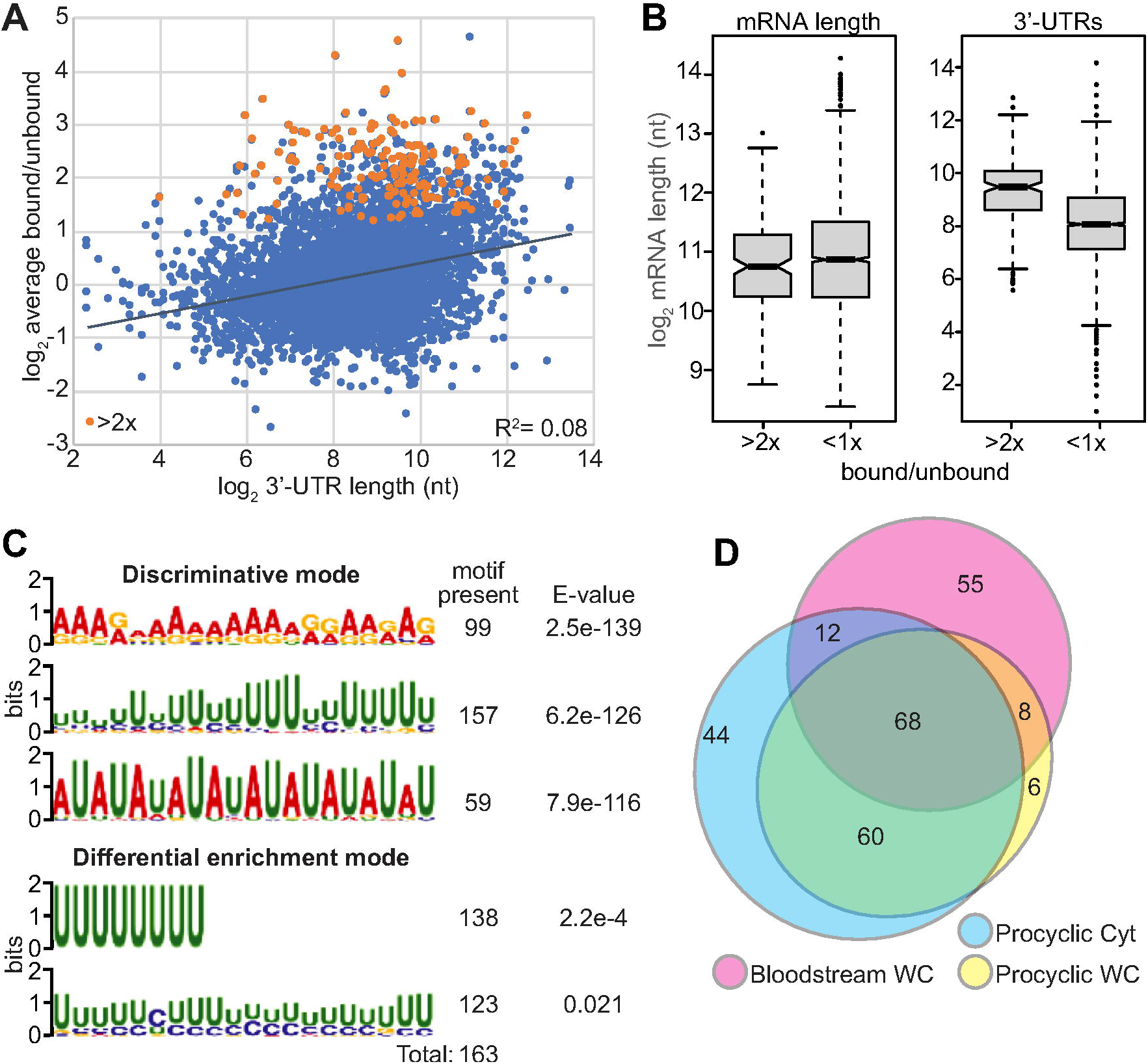
DRBD18 RNA binding and effects on the transcriptome. **A.** Tagged DRBD18 was pulled down from bloodstream-form extracts, then associated (bound) RNAs were released using TEV protease. RNA that did not bind to the affinity beads served as the negative control (“unbound”). The average bound/unbound ratio for three biological replicates is on the y axis and the annotated 3’-UTR length on the x-axis, both on a log scale. Each spots represents one open reading frame, with just one representative of repeated genes. The orange spots are genes with at least 10 reads and in all samples and with at least 2-fold enrichment in all three comparisons. **B.** Annotated mRNA lengths and 3’-UTR lengths for bound mRNAs (at least 2-fold enrichment in all three comparisons) and for mRNAs with, on average, less than 1-fold enrichment are plotted. **C.** The 3’-UTRs of the 163 bound mRNAs were compared with those of a length-matched set of mRNAs with less than 1-fold enrichment using MEME, using the discriminative mode and the more stringent differential enrichment modes. to the right are the numbers of bound mRNAs that contained the motif, and the probabilities from the programme. **D.** The log2fold change for each coding region was subtracted from the log2fold change of the corresponding 3’-UTR. If the result was less than −0.584 (1.5-fold excess of coding region reads) the mRNA was identified as having potential loss of 3’-UTR reads due to alternative polyadenylation. The numbers of genes for all three analysed conditions are shown to scale: Bloodstream-form whole-cell (WC) RNA, rRNA-depleted; Procyclic-form whole cell RNA, poly(A)+; and procyclic-form cytoplasmic RNA (Cyt), poly(A)+.

These results do not show that DRBD18 binds to PPTs - only that PPT motifs are enriched in its target mRNAs. Moreover, PPTs by themselves are clearly insufficient to distinguish between bound and unbound mRNAs. For example, there are 528 copies of (U)_8_, located in 138 of the 163 “bound” mRNA 3’-UTRs, but the 163 control “unbound” mRNAs also contain 157 copies of the same motif, located in 76 mRNAs and with up to 7 copies per 3’UTR.

### Effects of DRBD18 depletion on mRNA processing

To look for transcriptome-wide effects of DRBD18 depletion on mRNA processing, we compared the effects of RNAi on 3’-UTR reads with the effects on the coding region reads. The results for the 3’-UTRs are not as reliable as those for the coding-regions for two reasons. The 3’-UTRs contain many low-complexity sequences which may be repeated elsewhere in the genome, and more importantly, polyadenylation site mapping in the database is unreliable: many genes have no annotated 3’-UTRs, and the annotated ones are quite often shorter than those suggested by visual examination of the read densities. This means that some effects of DRBD18 depletion will have been missed. Nevertheless, for both bloodstream forms and procyclic forms, some mRNAs showed clear discrepancies between coding region and 3’-UTR reads (Supplementary Figure S3E, Supplementary Figure S4A), and the differences were especially marked when procyclic-form cytoplasmic poly(A)+ RNAs were examined (Supplementary Figure S3B). To find mRNAs with truncated 3’-UTRs after RNAi we chose those for which the change in 3’-UTR reads was at least 1.5-fold lower than the change in the coding region reads. This was true for 144 bloodstream-form whole-cell mRNAs, 142 procyclic-form whole-cell mRNAs, and 184 procyclic-form cytoplasmic mRNAs (Supplementary Table S5, sheet 2). There was a substantial overlap in these lists (Figure 5D, Supplementary Figure S4E). After elimination of obviously repeated genes, there were 253 coding regions. For bloodstream forms, 65 of the affected mRNA coding regions increased significantly in read density after RNAi and 17 decreased, while for procyclic forms, 99 increased and none decreased. Thus shortening of the mRNAs often, but not always, led to increases in abundance.

To analyse alternative processing in detail, we focussed on genes that showed a relative decrease in 3’-UTR reads in both procyclic and bloodstream forms (Supplementary Table S5, sheet 1), and for which the 3’-UTR had a single alternative processing site; the latter condition made quantitation possible. An example, Tb927.8.1580, is shown in Figure 6. In the absence of tetracycline reads were fairly uniformly distributed across the mRNA, the 3’-UTR of which is 1kb long (Figure 6A, Supplementary text 2). After RNAi induction, the read density over the coding region increased (Figure 6A, B), but there was an abrupt drop about 330nt into the 3’-UTR. resulting in a read density that was lower than in the presence of DRBD18 (Figure 6A). Scrutiny of the sequence reads (Figure 6C, Methods section and Supplementary Figure S5) revealed three new polyadenylation sites over a 40nt region that is about 90nt upstream of a 24nt PPT, which is in turn 28nt upstream of a new splice acceptor site (Figure 6C, Supplementary text S2, S3). The reads from downstream of the new poly(A) sites represent either full-length mRNA, or a 3’-UTR-derived RNA (shown as ** in Figure 6A). We counted reads from the new processing site region for all replicates (Figure 6C, D). It is important to note that we were analysing steady-state mRNA levels and we do not know the relative half-lives of the different Tb927.8.1580 RNAs. The results therefore do not quantitate the absolute frequency of alternative processing, but they do allow qualitative assessment of the effects of DRBD18 depletion. The top two bar-graph panels in Figure 6D show the numbers of reads covering the novel poly(A) sites, normalised to the total number of the reads in each dataset (Supplementary Table S5, sheet 6). The percentages of reads with alternative processing are shown as dot plots on the right. In the absence of tetracycline, all reads were wild-type (blue bars in Figure 6D). After RNAi, considering whole-cell RNA, about 30% of the reads that were captured for the two upstream internal poly(A)sites, p(A)-1/2, were polyadenylated there (orange bars in Figure 6D). 70% of the RNA would remain, and of these, about 30% terminated at poly(A) site 3. This analysis suggests that at least half of the mRNAs were alternatively processed at the 3’-end; the read density plots in Figure 6A suggest that the proportion was higher. Reads for the spliced leader acceptor site are shown in the lower part of Figure 6D. Only one read that showed this splicing event was observed in the presence of normal levels of DRBD18. After DRBD18 depletion, the number of full-length reads decreased, and 35% of the remaining reads were alternatively spliced. Interestingly, reads from alternatively spliced 3’-UTR-derived RNA were also present in the cytoplasmic fraction. The use of the alternative processing sites did not affect splicing of the downstream mRNA since there was no increase in intergenic reads (Figure 6A) and no reproducible effect on the abundance of the downstream mRNA, Tb927.8.1590 (Figure 6E).

**Figure 6:**
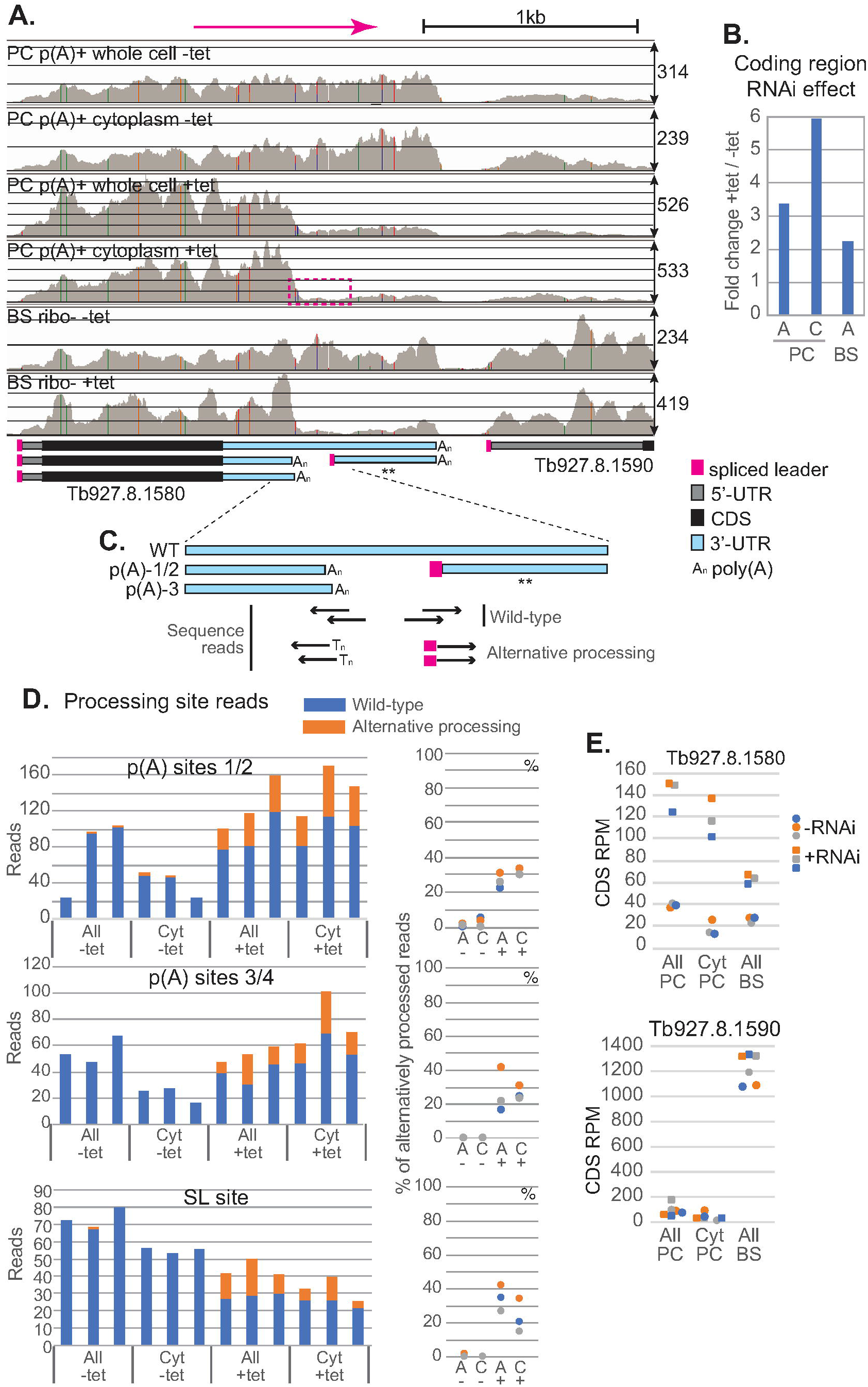
Effects of DRBD18 RNAi on the RNAs from Tb927.8.1580 and Tb927.8.1590. **A.** Results from the Integrated genome viewer in the region containing Tb927.8.1580 and Tb927.8.1590. The direction of transcription and a scale bar are at the top. Representative views for all six types of samples are shown. The scales on the y-axis are different (see numbers on the right); the thin lines are drawn at intervals that show read depths of approximately 100. A map of the detected mRNAs is below, as for Figure 1. **B.** Effects of RNAi on coding region reads (log2 fold change from DeSeq2). **C.** Method to estimate the relative abundances of wild-type and alternatively processed reads. For more detail, see Supplementary Figure S5. Reads that spanned the sites were designated as wild-type, and those with polyadenylation or spliced leader sequences were classified as alternatively processed. “All” - whole cell RNA; Cyt: cytoplasmic RNA **D.** The upper two bar graphs show numbers of reads (normalised to total dataset size) that were either wild-type of alternatively processed at the different poly(A) sites; all of these events are directed by *trans* splicing at a single downstream site, results for which are shown in the bottom panel. The dot plots show the proportions of mRNA that were alternatively processed (A=All). **E.** Reads per million for the coding regions of Tb927.8.1580 and the downstream gene, Tb927.8.1590, splicing of which directs wild-type Tb927.8.1580 polyadenylation. Results for all three replicates are shown for each dataset.

We repeated this analysis for 11 more genes. Examples of screen shots are in Supplementary Figures S6 and S7, and quantification of alternative site use is in Figure 7, Supplementary Figure S8, and Supplementary Table S5, sheet 6. The gene sequences with identified processing sites are in Supplementary text 2 and effects of RNAi on the abundances of the next downstream mRNAs are in Supplementary Figure S9. As expected from the read count values, the proportion of mRNAs that showed alternative polyadenylation was always higher after DRBD18 depletion. Usually, but not always, the shorter coding mRNA with alternative polyadenylation constituted a higher proportion of cytoplasmic RNA than the wild-type version; this could be because the shorter version was more readily exported, or had a longer cytoplasmic half-life, than the longer mRNA (e.g. Tb927.11.2500, Tb927.5.1130 and Tb927.7.18100 in Figure 7). Reads from the distal parts of the 3’-UTR were lower in the presence of tetracycline since we had specifically selected for this; and the short non-coding spliced products that resulted from use of 3’-UTR-internal PPTs had varying relative abundances in the cytoplasm. There was no reproducible effect of DRBD18 depletion on the abundance or splicing pattern of the downstream mRNA (Figure 7, Supplementary Figure S8, Supplementary Figure S9 and Supplementary Table S5, sheet 6). These observations suggest that at least for these mRNAs, loss of DRBD18 caused an increase in use of 3’-UTR-internal RNA processing signals, which was often accompanied by a decrease in the abundance of full-length mRNA. (We made a preliminary attempt to verify this for 5 mRNAs by Northern blotting, but the mRNAs are present at only about 1 copy per cell and were not detected.) DRBD18 appears to have no role in defining the next exon, since the coding mRNAs immediately downstream of alternatively polyadenylated mRNAs were not reproducibly affected.

**Figure 7:**
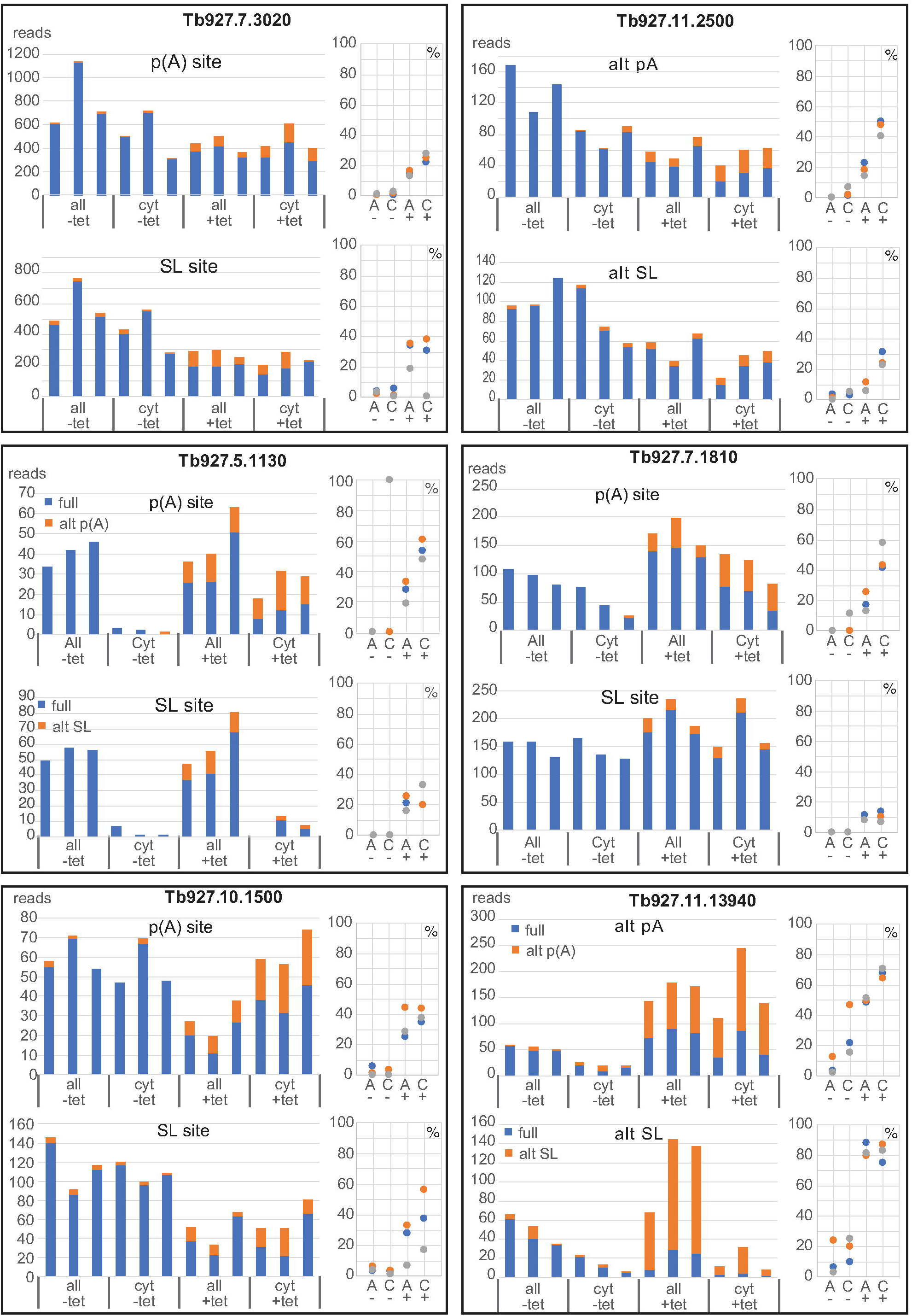
Effects of DRBD18 RNAi on additional mRNAs. For each gene, the upper two bar graph shows numbers of reads (normalised to total dataset size) that were either wild-type of alternatively processed at the different poly(A) sites, and the lower panel shows mRNAs with or without the 3’-UTR-internal splice site. Details are as in Figure 6.

**Figure 8.**
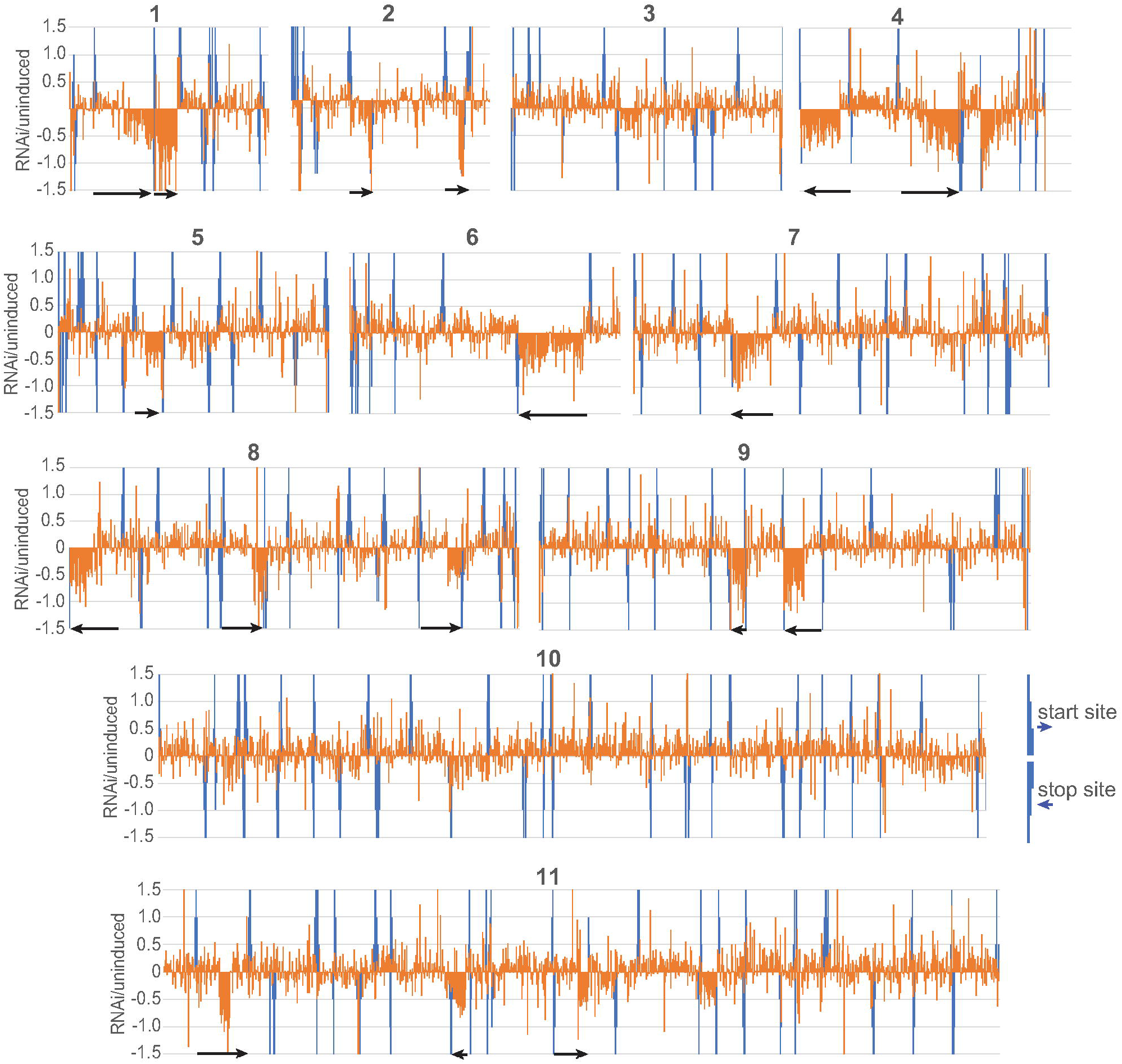
Effects of DRBD18 depletion show chromosomal clustering. The effects of DRBD18 depletion (log2 fold change) for unique genes (without paralogues) were plotted as orange bars according to the order of the genes on the chromosomes (positions not to scale). Start sites according to histone modifications (Siegel et al. 2009) and transcript orientation are upward blue bars, and stop sites are downward bars. Neighbouring smaller bars indicate the orientation. The scale of the y-axis has been truncated so that all effects greater than 2.8-fold are shown as log_2_(1.5). Black horizontal arrows indicate the directions of selected affected transcription units.

Finally, we looked at some of the mRNAs for which 3’-UTR reads increased, relative to coding-region reads, after DRBD18 RNAi (Supplementary Table S5, sheet 3). In some cases the result was an artefact due to low read coverage, or the presence of sequences that were found several times in the genome. In other cases, the 3’-UTR was annotated so as to overlap the 5’-UTR of the downstream gene, making the source of reads unclear and with no evidence for novel processing events. However, we also found instances that were indeed caused by alternative splicing events. In these cases, the novel non-coding RNA that arose from within the 3’-UTR was more abundant than the new truncated mRNA containing the coding region. One example, Tb927.6.2830, is shown in Supplementary Figure S7.

Interestingly, after RNAi in procyclic forms the 215 mRNAs that showed possible nuclear retention (or decreased stability in the cytoplasm) were significantly longer than the remainder (Supplementary Figure S4E, for definition see legend), and only five of them showed a possible change in processing. In contrast, half of the mRNAs with alternative processing showed an apparent increase in the ratio of cytoplasmic RNA to whole cell RNA after RNAi, suggesting that the new, shorter mRNAs were either more readily exported, and/or had increased stability in the cytoplasm relative to the longer species (Supplementary Table S5, sheet 2).

### Characteristics of alternatively processed mRNAs

We had 11 mRNAs for which we had mapped unique alternative processing sites and verified normal “wild-type” sites. This enabled us to investigate the difference between normal and novel sites. We considered two main possibilities. One possibility is that DRBD18 inhibits alternative processing by binding specifically upstream or downstream of the alternative splice sites; the other is that DRBD18 can bind all splice acceptor signals, but that at the wild-type splice site, DRBD18 effects are inhibited of other sequence-specific proteins. We therefore compared sequences from −50nt relative to the poly(A) sites to +50 downstream of the splice acceptor sites (Supplementary text S3). In pair-wise comparisons, there were no reproducible differences between the wild-type and alternative sites in PPT length or composition, distance to the acceptor AG, poly(A)-polypyrimidine distance, or GC content downstream of the splice site. There were also no differences in the numbers of additional intron PPTs that did not direct splicing. Splicing and transcription elongation kinetics can be influenced by RNA folding (see e.g. (Turowski et al. 2020)) but predicted folding energies of these RNA segments also showed no consistent differences. Finally, a comparison using MEME revealed no motifs that were specific to either the normal or the alternative sites. The sequences that differentiate wild-type from abnormal splice sites thus remain obscure. However, if wild-type sites are defined by various different proteins, with different specificities, or with binding sites that are more than 50nt downstream, we would not have detected the relevant sequence motifs with this small sample.

### DRBD18 depletion has polar effects on polycistronic transcription units

While examining the RNAi results, we noticed that some of the down-regulated genes appeared to be in clusters on the chromosomes. Most clusters were towards the ends of transcription units, but most transcription units were not affected (Figure 9). Similar results were seen for procyclic forms (not shown). Manual examination of the affected regions did not reveal any common features: in particular, mRNAs with obviously altered processing were only sometimes at the start of a repressed region. RNAi did not cause significant decreases in any mRNAs encoding known proteins involved in transcription. These results might therefore indicate hitherto unsuspected links between mRNA processing or export on transcription elongation.

**Figure 9.**
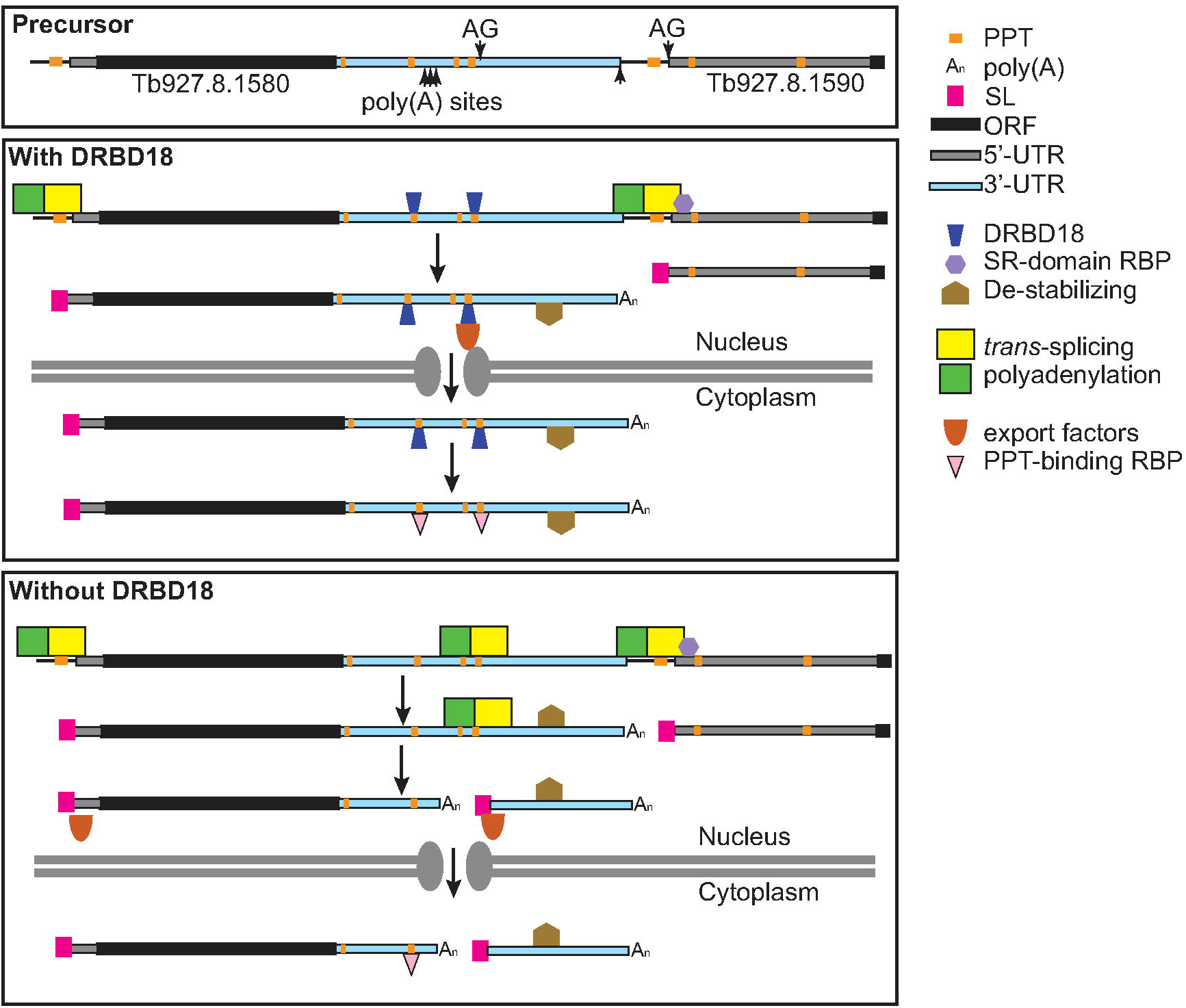
Model for the interaction between DRBD18 and *RBP10* mRNA. A key is on the Figure, which is explained in the text. RBP: RNA-binding protein.

## Discussion

The mechanisms by which splice acceptor sites are chosen in animal cells have been investigated extensively. As in trypanosomes, the presence of a polypyrimidine tract is insufficient to define a splice acceptor site. Instead, wild-type exons are marked by the binding of specific proteins (Ule and Blencowe 2019). For example, proteins that contain serine-arginine rich regions (SR domain) as well as an RNA recognition motif (RRM), bind to exonic sequences downstream of active splice sites (Howard and Sanford 2015). In trypanosomes, the only splicing signal that has been investigated in detail is the PPT. Results from one set of reporter experiments indicated that maximal splicing efficiency was obtained with PPTs of (U)_20_ or with 17mers of poly(U) containing interpelated C residues (Siegel et al. 2005), but another series indicated that a 10mer PPT interrupted by a single “A” was as good as a 20mer (Matthews et al. 1994). In practice, most mRNAs are *trans* spliced downstream of PPTs that are 12-24nt long, with more U than C (Kolev et al. 2010); but for some acceptors the PPT is barely recognisable. (Tb927.11.5450 and Tb927.11.5460 in Supplementary Text S3 are examples.) Excellent PPTs are scattered throughout 3’-UTRs, but are used only if the normal splice site has been removed (see e.g. (Hug et al. 1994; Matthews et al. 1994; Schürch et al. 1994; Vassella et al. 1994)). The most likely explanation is that splice sites that are upstream of coding regions are specifically chosen, as in animals; there is some extremely limited evidence for this from reporter experiments (Matthews et al. 1994; Hartmann et al. 1998).

We here show that DRBD18 preferentially binds mRNAs that contain several U-rich PPT motifs in their 3’-UTRs. Depletion of DRBD18 results in the accumulation of truncated mRNAs that arise from use of alternative splicing signals located in 3’-UTRs. In the examples examined, there was no evidence that DRBD18 loss had any effect on use of the normal splicing signal for the downstream mRNA. DRBD18 was already implicated in mRNA export: it interacts directly with MTR2, pulls down MEX67 (Mishra et al. 2021), and its depletion caused retention of poly(A)+ mRNA, MTR2 and MEX67 in the nucleus. We confirmed the association of DRBD18-containing mRNPs with proteins implicated in mRNA export and with the nuclear pore. Moreover, our results with bloodstream forms showed that the 8 kb wild-type *RBP10* mRNA was not exported from the nucleus in the absence of DRBD18. How can all of these results be interpreted? We considered several hypotheses.

The first possibility is that DRBD18 is required to define the position of the normal splice site. But if this were true, loss of DRBD18 should affect the abundance of the downstream mRNA or cause accumulation of RNAs that span the intergenic region. Neither effect was observed.

Another possibility is that alternative processing is constitutive, but the mRNAs with short 3’-UTRs, as well as the corresponding downstream spliced 3’-UTR fragments, are usually very unstable in the cytoplasm due to binding by DRBD18. If this were true, however, DRBD18 ought to destabilise the long mRNA as well, giving equal abundances of all products. To presere dominance of the longest mRNA it would be necessary to invoke some other protein that binds only to the long RNA - presumably just upstream of PPTs - and counteracts destabilizing effects or DRBD18. The hypothesis also fails to explain why in some cases, after DRBD18 depletion the abundance of full-length mRNAs decreases in favour of shorter variants.

A third idea might be that DRBD18 is required to export the longer mRNAs or stabilize them in the cytoplasm - but although neither effect be ruled out, they don’t explain why the shorter mRNAs only appear after DRBD18 loss. Moreover, the lack of correlation between steady-state DRBD18 binding and effects on mRNA abundance or processing - together with the preferential association of at least some DRBD18 mRNPs with the outer ring of the nuclear pore - suggest that after mRNA export into the cytoplasm is completed, mRNA-bound DRBD18 might be replaced by abundant PPT-binding proteins. Possible candidates might be the poly(U)-binding proteins UBP1 and UBP2 (D’Orso and Frasch 2001; Hartmann et al. 2007).

The hypothesis that we find to be most consistent with the data is shown in Figure 9. We speculate that in the nucleus, DRBD18 binds to PPTs in the 3’-UTRs of affected mRNAs. The previously-demonstrated accumulation of poly(A)+RNA in the nucleus after RNAi (Mishra et al. 2021) suggests that in some cases DRBD18 binding is required for (or stimulates) mRNA export. However, at least for *RBP10* mRNA, the effects of DRBD18 depletion on splicing and polyadenylation cannot solely be due to decreased export, because MEX67 depletion did not result in alternative processing. We therefore suggest that the binding of DRBD18 to 3’-UTR PPTs prevents their recognition by the splicing machinery, while perhaps also simultaneously promoting mRNA export. When DRBD18 is depleted, the newly-exposed alternative sites can be used. Truncated mRNAs that do not require DRBD18 for export may become abundant in the cytoplasm because they have lost destabilising 3’-UTR motifs that were present in the longer, wild-type version.

We do not know why the processing of some mRNAs is affected by DRBD18 loss, while others are unaffected, since PPTs are ubiquitously present in 3’-UTRs. We also do not yet know why wild-type splice sites are not inhibited by DRBD18, since in an (admittedly very small) sample no obvious sequence differences between them and the alternative sites were found. In humans, the sequences bound by different RNA-binding proteins often overlap, and it has been suggested that the context of a target sequence, or the secondary structure in the vicinity, is important in determining the extent of binding of particular proteins (Dominguez et al. 2018). The results of a recent study of a mouse RNA-binding protein indicated that its association and dissociation kinetics were affected by sequence context, and that association was enhanced if several binding sites were clustered (Sharma et al. 2021). The same may also apply to DRBD18: binding to mRNAs may be influenced by PPT frequency and context as well as competition with other proteins. The effects of DRBD18 could also be modulated by other proteins that bind the same mRNA precursor. By analogy with results from animal cells, DRBD18 activity at wild-type splice acceptor sites may be inhibited by proteins that show higher-affinity binding and specifically define exons. There is evidence for mRNA processing regulation by several nuclear trypanosome RNA-binding proteins (Stern et al. 2009; Gupta et al. 2013a; Gupta et al. 2013b; Gupta et al. 2014). Further analysis of these and other nuclear RNA-binding proteins will be needed to understand how sites for trypanosome RNA processing are normally determined.

## Materials and methods

### Trypanosome culture

The experiments in this study were carried out using monomorphic *T. brucei* Lister 427 (Alibu et al. 2004) expressing the tetracycline repressor. The bloodstream-form parasites were cultured as routinely in HMI-9 medium supplemented with 10% heat inactivated foetal bovine serum at 37°C with 5% CO_2_. During proliferation, the cells were diluted to 1×10^5^ cells/ml and maintained in density between 0.2-1.5×10^6^ (Clayton et al. 2005). For generation of stable cell lines, ~1-2 x 10^7^ cells were transfected by electroporation with 10 μg of linearized plasmid at 1.5 kV on an AMAXA Nucleofector. Selection of newly transfectants was done after addition of appropriate antibiotic and serial dilution. The induction of RNAi was done using 100 ng/ml tetracycline, in the absence of other selective antibiotics.

### Plasmid constructs

For endogenous tagging of DRBD18, a cell line with *in-situ* TAP-DRBD18 was generated by replacing one endogenous copy of DRBD18 with a gene encoding a C-terminally TAP-tagged DRBD18. For that, a construct with neomycin resistance gene plus TAP tag cassette was flanked on the 3’-end with a fragment of *DRBD18* 3’-UTR. Upstream on the 5’-end, the C-terminal region of the *DRBD18* ORF without the stop codon was cloned in frame with the TAP tag. Prior to transfection in monomorphic bloodstream forms Lister 427, the plasmid (pHD3200) was cut with *Apa* I and *Xba* I to allow homologous recombination. The tetracycline inducible construct for DRBD18 RNAi was done using a stem-loop vector targeting the coding region of DRBD18. The region used for the stem loop vector was checked against off-targets using the RNAi target selection tool for trypanosome genomes (https://dag.compbio.dundee.ac.uk/RNAit/) as described in (Redmond et al. 2003). Plasmids and oligonucleotides are listed in Supplementary Table S7.

### RNA analysis and Northern Blotting

Total RNA was isolated from approximately 1×10^8^ bloodstream-form trypanosome cells using either peqGold Trifast (PeqLab) or RNAzol RT following the manufacturer’s instructions. To detect the mRNA by Northern blot, 5 or 10 μg of the purified RNA was resolved on formaldehyde agarose gels and transferred onto nylon membranes (GE Healthcare) by capillary blotting and fixed by UV-crosslinking. The membranes were pre-hybridized in 5x SSC, 0,5 % SDS with 200 mg/ml of salmon sperm DNA (200 mg/ml) and 1x Denhardt’s solution, for an hour at 65°C. The probes were generated by PCR of the coding sequences of the targeted mRNAs, followed by incorporation of radiolabelled [α^32^P]-dCTP and purification using the QIAGEN nucleotide removal kit according to the manufacturer’s instructions. The purified probes were then added to the prehybridization solution and the membranes were hybridized with the respective probes at 65°C for overnight (while rotating). After rinsing the membranes in 2x SSC buffer/0.5% SDS twice for 15 minutes, the probes were washed out once with 1x SSC buffer/0.5% SDS at 65°C for 15 minutes and twice in 0.1x SSC buffer/0.5% SDS at 65°C each for 10 minutes. The blots were then exposed onto autoradiography films for 24-48 hours and the signals were detected with the phosphorimager (Fuji, FLA-7000, GE Healthcare). The signal intensities of the images were measured using ImageJ.

Total RNA used for sequencing analysis (RNA-Seq) was prepared from cells collected 12 hours after the induction of DRBD18 RNAi in monomorphic bloodstream forms cells (Lister 427). Induction of RNAi was done using tetracycline at a concentration of 100 ng/ml. The cells without induction of DRBD18 RNAi were used as controls. RNA was extracted using the phenol-chloroform separation method as described previously. The integrity of the total RNA was checked on a denaturing agarose gel. Afterwards, the ribosomal RNAs (rRNAs) were depleted from the total RNA samples using a cocktail of 131 50-base oligonucleotides complementary to rRNAs, combined with RNaseH (NEB, M0297S), as previously described (Antwi et al. 2016; Minia et al. 2016). Following rRNA depletion, the samples were subjected to DNAse I treatment in order to remove oligonucleotides using the Turbo DNAse kit (Invitrogen, ThermoScientific). The RNA samples were then purified using the RNA Clean & Concentrator −5 kit (ZYMO RESEARCH) following the manufacturer’s instructions. The recovered purified RNA from both bound and unbound samples was then analysed by RNA-Seq..

### RNA immunoprecipitation

A cell line expressing *in-situ* C-terminally TAP tagged DRBD18 was used for the RNA immunoprecipitation. The TAP-tag consists of the protein A and calmodulin binding protein domains separated by a Tobacco Etch Virus (TEV) protease cleavage site. Approximately 1×10^9^ cells expressing *in-situ* C-TAP DRBD18 with a concentration of 1×10^6^ cells/ml were pelleted by centrifugation at 3000 rpm for 13 minutes at 4°C. The pellet was washed twice in cold 1x PBS and collected by centrifugation at 2300 rpm for 8 minutes at 4°C and then snap frozen in liquid nitrogen. The cell pellet was lysed in 1 ml of the lysis buffer (20 mM Tris pH 7.5, 5 mM MgCL_2_, 0.1% IGEPAL, 1 mM DTT, 100 U RNAsin, 10 μg/ml leupeptin, 10 μg/ml Aprotinin) by passing 20 times through a 21G x ½ needle using a 1 ml syringe and 20 times through a 27G x ¾ needle using a 1 ml syringe. The lysate was cleared by centrifugation at 15,000 g for 15 minutes at 4°C. Afterwards, the supernatant was transferred to a new Eppendorf tube and the salt concentration was adjusted to 150 mM KCl. 10% of lysate was collected as the input fraction for RNA extraction and 2% was used for western blotting. 40 μl of magnetic beads (Dynabeads M-280 Tosyl activated, Invitrogen) coupled with Rabbit Gamma globulin antibodies (Jackson Immuno Research Laboratories) were washed three times with 500μl of IP buffer (20 mM Tris pH 7.5, 5 mM MgCL_2_, 150 mM KCl, 0.1% IGEPAL, 1 mM DTT, 100 U RNAsin, 10 μg/ml leupeptin, 10 μg/ml Aprotinin) then incubated with the cell lysate at 4°C for 3 hours with gentle rocking. The unbound fraction was collected for RNA extraction (98%) and Western blotting (2%). Subsequently, the beads were washed thrice with IP buffer. 2% of each wash fraction was collected for protein detection. The TAP-tag was cleaved by incubating the beads with 500 μl of IP buffer containing 100 units of TEV protease (5 μl) with gentle rotation at 16°C for 2 hours. The eluate was then collected afterward by magnetic separation for RNA isolation and protein detection. RNA was isolated from the input, the unbound and the eluate fraction using the peqGOLD Trifast FL (Peqlab, GMBH) according the manufacturer’s instructions. To assess the quality of the purified RNA, aliquots of the input, unbound and eluate fractions were resolved on formaldehyde agarose gels to check the integrity of the ribosomal RNAs.

Ribosomal RNAs were then depleted as described above and the fractions sent for RNA sequencing.

### High throughput RNA sequencing and bioinformatic analysis

RNA Sequencing was performed at the Cell Networks Deep Sequencing Core Facility of the University of Heidelberg. The library preparation was done using the NEBNext Ultra II Directional RNA Library Prep Kit for Illumina (NEB, E7760S). The libraries were multiplexed (six samples per lane) and sequenced with a NextSeq 550 system, generating single-end sequencing reads of about 75 bp. Raw data are available at Annotare as E-MTAB-9783 and E-MTAB-10735.

Sequence alignment and read counting were done using custom pipelines (Leiss and Clayton 2016; Leiss et al. 2016). Before analysis, the quality of the raw sequencing data was checked using FastQC (http://www.bioinformatics.babraham.ac.uk/projects/fastqc). Cutadapt (Martin 2011) was used to remove sequencing primers, then the sequencing data were aligned to *T. brucei* 927 reference genome using Bowtie (Langmead and Salzberg 2012) allowing 1 alignment per read, then sorted and indexed using SAMtools (Li et al. 2009). For RNAi, the reads that mapped to the open reading frames, 3’-UTRs and non-coding RNAs in the TREU 927 genome were counted. Alignment and read-counting for the published results for procyclic-form whole-cell and cytoplasmic RNA (Mishra et al. 2021) was done using the same pipelines. Reads per million reads (RPM) were calculated after removal of rRNA reads, and the ratios of bound RPM to unbound RPM were calculated for each of the three purifications. To find reproducibly bound mRNAs, we took into account only the lowest ratio for each mRNA.

In order to look for enrichment of particular functional characteristics, we used a list of unique genes modified from (Siegel et al. 2010); this corrects for repeated genes and multigene families. To generate reads per million reads for the unique gene set, we first multiplied the reads for each gene in the list by the gene copy number (obtained in a separate analysis, see e.g. (Mulindwa et al. 2021)). For the RIP-Seq, the reads per million reads were counted and the ratios of eluate versus flow-through were calculated. An mRNA was considered as “bound mRNA” if the ratios from all the three pulldowns were higher than 1.5. The 3’-UTR motif enrichment search was done using MEME in the relative enrichment mode (Bailey 2011). Annotated 3’-UTRs were downloaded from TritrypDB. For some other 3’-UTRs, the manual annotation was performed using the RNA-Seq reads (e.g. (Jensen et al. 2014) and poly(A) site data in TritrypDB (Kolev et al. 2010). Analysis of differentially expressed genes after DRBD18 RNAi was done in R using DESeqUI (Leiss and Clayton 2016), a customized version of DESeq2 (Love et al. 2014) adapted for trypanosome transcriptomes. Statistical analyses were done using R and Microsoft excel.

To find alternative processing sites, aligned reads were visualised using the Integrated Genome Viewer (Robinson et al. 2011; Thorvaldsdóttir et al. 2013). Terminal mismatches suggesting the presence of poly(A) tails or spliced leaders were identified. We then selected 17-20nt sequences immediately upstream of the putative poly(A) sites or immediately downstream of the splice sites, to extract all overlapping reads from the raw data. This enabled us to find reads that had too many terminal mismatches to be present in the final aligned files. We used only the procyclic-form poly(A)+ results for this; our bloodstream-form reads had almost no spliced leaders of poly(A) tails, perhaps due to the RNase H treatment. The lengths of sequences used for the extraction, and their distances from the processing sites, were not always the same, because highly repetitive sequences could not be used. Therefore, the proportion of overlapping reads extracted is expected to differ for different sites. The extracted reads were then edited manually to remove reads that come from the wrong gene, or had insufficient terminal bases to identify processing. Manual examination was particularly important for polyadenylation since often, several different alternative poly(A) sites were found within the region upstream of the new splice site. The numbers of sequences that spanned the processing site, or were processed, were then counted for all replicates.

RNA folding was predicted using the RNAFold Web server (http://rna.tbi.univie.ac.at/cgi-bin/RNAWebSuite/RNAfold.cgi).

### Western blotting

Protein samples were collected from approximately 5×10^6^ cells growing at logarithmic phase. Samples were run according to standard protein separation procedures using SDS-PAGE. The primary antibodies used in this study were: rabbit α-DRBD18 (1:2500) (Lott et al. 2015), rat α-ribosomal protein S9 (unpublished) and rabbit anti-XRND (1:2500 (Li et al. 2006)). We used horseradish peroxidase coupled secondary antibodies (α-rat, 1:2000 and α-rabbit, 1:2500). The blots were developed using an enhanced chemiluminescence kit (Amersham) according to the manufacturer’s instructions. The signal intensities of the images were quantified using ImageJ.

### Protein-protein interactions

A cell line in which one allele of DRBD18 bears a sequence encoding a C-terminal TAP tag was used to study the interactome of DRBD18. DRBD18-TAP was purified using IgG magnetic beads and the bound protein was eluted with TEV protease, which was then removed using HisPur Ni-NTA magnetic beads (Thermo Scientific) according to the manufacturer’s instructions (Falk et al. 2021). The proteins from three independent purifications were subjected to denaturing SDS-PAGE and analysed by mass spectrometry. A cell line inducibly expressing GFP with a TAP tag at the N-terminus was used as a control. The expression of GFP-TAP was induced with 100 ng/ml tetracycline for 24 hours and purification of the protein was done as for DRBD18-TAP. After purification, the proteins were subjected to denaturing SDS-PAGE, visualized by Coomassie blue staining and analysed by mass spectrometry (Chakraborty and Clayton 2018). Data were quantitatively analysed using Perseus (Tyanova et al. 2016).

### Subcellular fractionation

Nuclear-cytosolic fractionation was performed as described in (Biton et al. 2006). Briefly, 3×10^8^ Lister 427 bloodstream form wild-type cells as well as those expressing the RNAi stem loop vector targeting DRBD18 were harvested by centrifugation at 3000 rpm for 13 minutes at 4°C. The cell pellet was washed twice with cold 1x PBS and collected by centrifugation at 2300 rpm for 8 minutes at 4°C. Afterwards, the pellet was resuspended in hypotonic buffer (10 mM HEPES pH 7.9, 1.5 mM MgCl2, 10 mM KCl, 0.5 mM dithiothreitol, 5 μg/ml leupeptin and 100 U RNasin) and lysed in presence of 0,1% IGEPAL (Nonidet P-40) by passing 20 times through a 21G x ½ needle using a 1 ml syringe and 20 times through a 27G x ¾ needle using a 1 ml syringe. The cells were allowed to rest on ice for 20 minutes. Then, the nuclei fraction was pelleted by centrifugation at 15,000 g for 15 minutes at 4°C. The supernatant containing the cytoplasmic fraction was transferred to a new tube. 20 % of the cytoplasmic fraction was used for western blotting and the rest was used for RNA extraction. The pellet containing the nuclei fraction was resuspended in TBS containing 0.1 % SDS; 20 % of this fraction was used for western blotting while the rest was used for RNA extraction (result not shown).

### Affymetrix smRNA FISH

The Affymetrix single molecule mRNA FISH was carried out as described in (Kramer 2017). A total of 200 ml bloodstream-form trypanosomes at ~ 5-8 x10^5^ cells/ml were harvested by centrifugation (8 minutes, 1400g), resuspended in 1 ml 1x PBS and pelleted again by centrifugation (5 minutes, 1400g). The cell pellet was resuspended in 1 ml 1x PBS, followed by fixation with 1 ml of 8% formaldehyde (in PBS). The mixture was incubated at room temperature for 10 minutes with an orbital mixer. A total of 13 ml 1x PBS were added and the cells were harvested by centrifugation (5 minutes, 1400 g). The pellet of fixed cells was resuspended in 1 ml 1x PBS and spread on glass microscopy slides (previously incubated at 180°C for 2h for RNase removal) within circles of hydrophobic barriers (PAP pen, Sigma). The cells were allowed to settle at room temperature for 20 minutes. The slides were then washed twice in 1x PBS. The protease solution was diluted 1:1000 in 1x PBS and briefly vortexed to allow complete dissolution. Permeabilization of the fixed cells was done with addition of 50 μl of detergent solution QC in each circle on the slides. This was followed by a 2-step washing in 1x PBS. 100 μl of the protease solution was added to each circle and incubated exactly for 10 minutes at 25°C. The slides were then washed twice in 1x PBS and used for Affymetrix FISH experiments as described in the manual of the QuantiGene ViewRNA ISH Cell Assay (Affymetrix), protocol for glass slide format. The only modification from the kit protocol is that the protease digestion was done at 25°C rather than the normal room temperature and we used a self-made washing buffer (0.1x SSC buffer, 0,1% SDS) instead of the washing buffer from the kit. All Affymetrix probe sets used in this work are described in Supplementary Table S7 (sheet 3). For visualization, the labelled cells were mounted with 4′,6-diamino-2-phenylindole dihydrochloride (DAPI) solution, diluted 1:1000 in 1x PBS. Images were taken with a fluorescent inverted wide-field microscope Leica DMI6000B (Leica Microsystems GmbH, Wetzlar, Germany) equipped with 100x oil immersion (NA 1.4) and a Leica DFC365 camera (6.45 m/pixel). Deconvolution was done using Huygens Essential software (SVI, Hilversum, The Netherlands) and images are presented as Z-stack projection (sum slices). The image analysis was carried out using the available tools in Image J software and 50 cells for each slide were selected for quantifying the number of mRNAs present in the cytosol and in the nucleus.

## Acknowledgments

We are very grateful to Dr. Susanne Kramer, who supervised and provided facilities for the mRNA FISH experiment. We are thankful to Prof. Dr. Laurie Read for antibody donation and for extensive and friendly discussions, including communicating unpublished results. We thank Bernardo Gabiatti and Paula Andrea Castañeda Londoño (Biozentrum, Universität Würzburg) for discussion and assistance with the mRNA FISH experiment; Laura Armbruster (COS, University of Heidelberg) for help in mass spectrometry data analysis, and Kevin Leiss, Abeer Fadda and Simon Anders for supplementing CC’s rudimentary bioinformatics skills. All RNA-Seq libraries and RNA sequencing were done by David Ibberson at the Bioquant in the University of Heidelberg. Mass spectrometry was done in the ZMBH Core Facility for Mass Spectrometry by Thomas Ruppert and Sabine Merker. We thank Claudia Helbig and Ute Leibfried for technical assistance, for preparing media and buffers. We are indebted to Prof. Dr. Nina Papavasiliou (DKFZ, University of Heidelberg) and Prof. Dr. Luise Krauth-Siegel (BZH, University of Heidelberg) for allowing us to share their laboratories including equipment and reagents after a flood in ZMBH. This work was partially supported by the Deutsche Forschungsgemeinschaft, grant number Cl112/28-1 to CC, and partially by core funding.

## Data availability

Raw RNA-Seq data are available at Annotare as E-MTAB-9783 and E-MTAB-10735. The mass spectrometry proteomics data have been deposited to the ProteomeXchange Consortium via the PRIDE partner repository (Perez-Riverol et al. 2019) with the dataset identifier PXD027792.

## Author contributions

Tania Bishola Tshitenge was responsible for nearly all of the experimental work, provided figures and tables, wrote the first draft of the paper and was involved in subsequent editing. Christine Clayton devised and supervised the project, analysed the procyclic-form RNA-Seq data, edited the paper and provided funding.

